# From the lung to the muscle: Systemic insights from an integrative MultiOmics analysis of harbour porpoises in poor respiratory health

**DOI:** 10.64898/2026.03.28.714973

**Authors:** Eda Merve Dönmez, Bente Siebels, Bernhard Drotleff, Paula Nissen, Davina Derous, Andrej Fabrizius, Ursula Siebert

## Abstract

Harbour porpoises (*Phocoena phocoena*) in the North and Baltic Seas are increasingly impacted by anthropogenic pressures, including underwater noise, fisheries and pollution. These pressures correlate with declining population health, particularly affecting the respiratory system. Growing pathological lesions, partly resulting from high prevalence of parasitic infestations and subsequent diseases, can impair tissue function and oxygen supply to distant end-organs.

In this study, we applied an integrative MultiOmics approach (proteomics, metabolomics, lipidomics) to analyse the lungs and muscles of 12 wild harbour porpoises with compromised respiratory health. Our aim was to identify dysregulated biological pathways across omics layers to advance insights into adaptive physiological responses and to define disease-associated molecular signatures that could assist health assessments.

Our analysis revealed pronounced immune system and antioxidative responses in the lungs and muscles, indicated by enhanced immunoglobulins, plasmalogens and glutathione-related proteins. In the lungs, high cardiolipin levels and reduced collagen suggest impaired tissue structure and function, while tissue maintenance processes were elevated in the muscle. Both tissues exhibited metabolic alterations suggestive of energetic imbalance, including increased purine metabolism in the lung and decreased lipid metabolism in the muscle. Several dysregulated molecules were shared across tissues, pointing to pathophysiological effects. The proposed disease-associated molecular signatures included the protein SLC25A4, the metabolite O-phosphoethanolamine and the lipid TG O-16:0_16:0_20:4 for the lung, and the protein SPEG, the metabolite pipecolic acid, and the lipid BMP 18:1_22:6 in the muscle.

Our findings elucidate the complexity of molecular mechanisms linking anthropogenic and environmental stressors with vulnerability and resilience in a marine sentinel species. Furthermore, this study highlights the potential of integrative omics to define disease-related marker panels, thereby supporting ongoing and future health monitoring and conservation efforts.

## INTRODUCTION

The harbour porpoise (*Phocoena phocoena*) is a cetacean abundant across the North Atlantic and frequently encountered in coastal areas. In the North and Baltic Seas, the harbour porpoise also functions as a sentinel species indicating ecosystem status, and several distinct subpopulations with differing habitats and conservation concerns exist (Carlström et al., 2023; Koschinski et al., 2024; Nachtsheim et al., 2021; Owen et al., 2024). As economic interests in these waters rise, their impact on cetaceans results in cumulative stress which can reduce fitness and cause higher morbidity (Carlén et al., 2021; Kesselring et al., 2017; Siebert et al., 1999; Vinther & Larsen, 2004; Wisniewska et al., 2018). Although many natural and anthropogenic pressures affect harbour porpoises, underwater-radiated noise is a growing concern as dependence on maritime infrastructure and construction continuously increases (Halpern et al., 2008). With several offshore constructions and important harbours located in German waters, noise pollution accumulates and interferes with the porpoises’ communication, hearing and behaviour (Erbe et al., 2019; Kastelein et al., 2018). Free-ranging individuals display a profound flight-and-avoidance reaction when encountering ships or acoustic deterrence signals, including spontaneous, for the species atypical prolonged and deeper dives (Elmegaard et al., 2023; Frankish et al., 2023; Wisniewska et al., 2018). While harbour porpoises generally perform short dives within 30-50 m, as the North and Baltic Seas are mostly shallow waters (Teilmann et al., 2007; Westgate et al., 1995), tagged porpoises were able to perform deeper dives, if the bathymetry allowed such dive profiles (Teilmann et al., 2007).

Diving demands large amounts of energy for the muscles to fuel force and speed, for which adequate oxygen supply is necessary. To enable diving bouts, marine mammal muscles possess higher myoglobin concentrations to store oxygen (Arregui et al., 2021), and are more tolerant of anaerobic metabolism to sustain energy levels upon oxygen depletion (Castellini et al., 1981). In addition to the blood and muscles, harbour porpoises also use their lungs for oxygen storage (Piscitelli et al., 2010). However, overall health and especially the pulmonary health of harbour porpoise populations is declining (Ryeng et al., 2022; Siebert et al., 2001, 2009). Their lungs are commonly found with infestations of nematodes that can cause local inflammation, obstruction of blood vessels and increase the susceptibility to additional diseases (IJsseldijk et al., 2021; Jepson et al., 2000; Siebert et al., 2001; Siebert et al., 2006a). Aberrant pathological lesions may not only obstruct the capacity of the lung to take up sufficient oxygen, but may also affect distant, oxygen-dependent organs, such as the muscles (Ceco et al., 2017; Siebert et al., 2001; Ten Doeschate et al., 2017). Inadequate oxygen supply to the locomotor muscles may result in impaired diving ability, if the disparity between muscular oxygen demand and supply by the lung is too large to compensate. In captive harbour porpoises, it has been observed that they reach only ∼60% of the complete lung capacity (Rojano-Donãte et al., 2018). However, it has yet to be clarified, whether the reduced utilisation is due to tissue function impairment or if it represents a voluntary behaviour and harbour porpoises do not need to exploit their full lung capacity.

Although many adaptations of marine mammals have been extensively investigated (Allen & Vázquez-Medina, 2019; Davis, 2014; Hindle, 2020), some molecular and physiological mechanisms underlying diving in cetaceans remain unresolved. Studies on cetaceans are limited by access to samples due to their fully aquatic lifestyle and conservation status (Van Cise et al., 2024). Additionally, reference databases of non-model organisms, such as marine mammals, are not always available and complete for various tools (Cammen et al., 2016), hence hinder in-depth analyses. Omics studies offer an alternative to gain knowledge about environmental impacts and molecular adaptations, even if sample sizes are small and limited. In marine mammal science, omics are an emerging tool and have been mostly applied on cetacean biopsy or blood samples (Kershaw et al., 2018, 2024; Morey et al., 2022; Trego et al., 2019; Veldhoen et al., 2012). Current studies often rely on one or two omics layers to analyse dysregulated biological processes and pathways (Derous et al., 2022; Houser et al., 2021; Simond et al., 2025; Trego et al., 2019). Combining multiple omics methods provides holistic insights into systems biology and cross-layer regulation, particularly for inaccessible, non-model organisms (Mancia, 2018). Moreover, a deeper understanding of molecular and physiological responses to disease and stress in cetaceans is crucial to assess population resilience and vulnerability under intensifying environmental pressures (Lennon et al., 2025), thereby supporting long-term monitoring efforts.

Here, we performed comparative, untargeted MultiOmics including proteomics, metabolomics and lipidomics of the lungs and locomotor muscles of free-ranging harbour porpoises that died from stranding or as by-catch. We compared porpoises suffering from severe pathological lung lesions, partly due to nematode infestations and bronchopneumonia, against porpoises without lung-associated diseases to assess dysregulated biological pathways. For both tissues, we further analysed the correlation between the health status and dysregulated features to tentatively identify a lung disease-associated, molecular signature that may serve as an analytical tool to advance disease diagnostics and support conservation efforts.

## MATERIALS AND METHODS

### Animals and Sampling

Tissue samples of the lung and muscle (*Musculus longissimus dorsalis*) were collected during routinely performed necropsies of harbour porpoises (n = 13, Tab. 1) at the Institute of Terrestrial and Aquatic Wildlife Research, University of Veterinary Medicine Hannover, Foundation, Büsum, Germany, which is part of the German stranding network (Benke et al., 1998; Siebert et al., 2006b). Harbour porpoises died by stranding or were by-caught between April 2016 and August 2022. Decomposition was not advanced (decomposition code II, estimated within 12-24 h of death). Full necropsies and further investigations of the animals were performed according to standardised protocols (IJsseldijk et al., 2019; Siebert et al., 2001). Depending on the necropsy findings, harbour porpoises were categorized as non-healthy or healthy. Harbour porpoises classified as non-healthy exhibited considerable pathological lesions due to nematode infestations (lungworms) and subsequent bacterial infections in the lung, and suffered or died from bronchopneumonia (Tab. 1). Healthy porpoises displayed minor to no pathological lesions in the lung and were in good overall health. Tissue samples were stored at -20°C until further use.

**Table 1.**
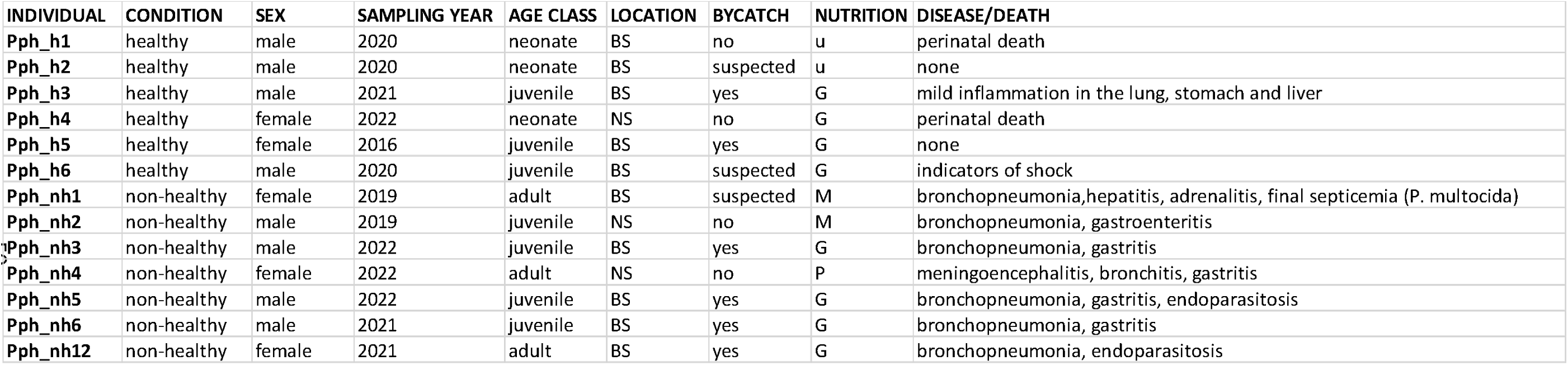
Metadata of the wild harbour porpoises used in this study. Health condition, sex, year of sampling, age class, habitat, bycatch, nutritional state and disease/cause of death are given. Non-healthy harbour porpoises were defined as animals that suffered from severe levels of lungworm infestation and bronchopneumonia. Healthy individuals did not suffer or die from lung-associated diseases or infestations. Nutritional state is defined as G: good, M: moderate, P: poor and u: undetermine d. Animals were predominatel y from the Baltic Sea (BS) and three individuals were from the North Sea (NS).

### LC-MS/MS-based bottom-up proteomics

#### Tryptic digest

For the proteomic analysis, 10 mg of frozen lung and muscle tissue were provided to the Core Facility Mass Spectrometric Proteomics of the University Medical Center Hamburg-Eppendorf.

Samples were dissolved in 100 mM triethyl ammonium bicarbonate and 1% w/v sodium deoxycholate buffer, boiled at 95°C for 5 min and sonicated with a probe sonicator. The protein concentration of denatured proteins was determined with the Pierce™ BCA Protein assay kit (Thermo Fisher, USA), and samples were diluted to 20 μg protein in 50 µL buffer. Samples were dissolved to a concentration of 70% acetonitrile (ACN). Carboxylate-modified magnetic beads (1 µL, GE Healthcare Sera-Mag™, Chicago, USA) at 1:1 (hydrophilic/hydrophobic) ratio in methanol/LC-MS-grade water were added, following the SP3-protocol workflow (Hughes et al., 2019). Samples were shaken at 1400 rpm for 18 min at room temperature. Tubes were placed on a magnetic rack and supernatant was removed. Magnetic beads were washed two times with 100% ACN and 70% ethanol. After resuspension in 50 mM ammonium bicarbonate, disulfide bonds were reduced in 10 mM dithiothreitol for 30 min and alkylated in presence of 20 mM iodoacetamide for 30 min in the dark. Samples were digested with sequencing-grade trypsin (Promega, USA) at 1:100 (enzyme:protein) ratio at 37°C and shaken at 1400 rpm overnight. Tryptic peptides were bound to the beads by adding 95% ACN and shaken at 1400 rpm for 10 min at room temperature. Placed on the magnetic rack, the supernatant was removed and the beads were washed twice with 100% ACN. Elution was performed with 2% DMSO in 1% formic acid (FA). The supernatant was dried in a vacuum centrifuge and stored at -20°C until further use.

#### LC-MS/MS data acquisition

Chromatographic separation of peptides was achieved with a two-buffer system (buffer A: 0.1% FA in H_2_O; buffer B: 0.1% FA in ACN) on a nano-Ultra High-Performance Liquid Chromatography system (Dionex Ultimate 3000 UPLC with a NanoViper C18 peptide trap (5 µm, 100 µm x 20 mm), Thermo Fisher, USA) for online desalting and purification, followed by a nanoEase M/Z Peptide BEH C18 column (1.7 µm, 75 µm X 250 mm; Waters, USA). Peptides were separated using an 80 min-method with linearly increasing ACN concentration from 2% to 30% over 60 min. MS/MS measurements were performed on a quadrupole-orbitrap hybrid mass spectrometer (QExactive, Thermo Fisher, USA). Eluting peptides were ionized using a nano-electrospray ionization source with a spray voltage of 1800 and analysed in data-dependent acquisition mode. For each MS1 scan, ions were accumulated for a maximum of 240 ms or until a charge density of 1 x 10^6^ ions (AGC target) was reached. Fourier-transformation-based mass analysis of the data from the orbitrap mass spectrometer was performed covering a mass range of 400 – 1200 m/z with a resolution of 70,000 at m/z = 200. Peptides, responsible for the 15 highest signal intensities per precursor scan with a minimum AGC target of 5 x 10^3^ and charge state from +2 to +5, were isolated within a 2 m/z isolation window and fragmented with a normalised collision energy of 25% using higher energy collisional dissociation. MS2 scanning covered a mass range starting at 100 m/z and was accumulated for 50 ms or to an AGC target of 1 x 10^5^ at a resolution of 17500 at m/z = 200. Already fragmented peptides were excluded for 20 s.

#### Data processing and quantification

LC-MS/MS data were searched with the Sequest algorithm, integrated into the Proteome Discoverer software (v 3.0; Thermo Fisher, USA), against a reviewed vaquita database (*Phocoena sinus*, TaxID=9741_and_subtaxonomies, 2023). Lung and muscle tissue samples were searched separately. Carbamidomethylation was set as a fixed modification for cysteine residues. The oxidation of methionine and pyro-glutamate formation at glutamine residues at the peptide N-terminus, as well as acetylation of the protein N-terminus were allowed as variable modifications. A maximum number of two missing tryptic cleavages was set. Peptides between 6 and 144 amino acids were considered. A strict cut-off (FDR < 0.01) was set for peptide and protein identification. Quantification was performed using the Minora algorithm, implemented in Proteome Discoverer, using the match-between-runs function.

#### Statistical analysis

Obtained protein abundances were log_2_-transformed and normalised by column-median normalisation. As the sample preparation was performed in two batches, batch effects between the two measurements were removed using the BERT algorithm (Schumann et al., 2025). In total, 2560 proteins were identified in lung tissue and 1526 proteins in muscle tissue. The data was reduced to proteins found in at least 70% of all samples and a two-sided unpaired Student’s T-test was performed using the statistic program Perseus (Tyanova et al., 2016), including permutation-based false discovery rate (FDR)-correction with 250 randomisations and q-value cut-off of 0.05.

One individual was not included in the subsequent analyses, since it was an outlier in multiple experiments. This may be due to advanced tissue degradation, possibly due to freeze-and-thawing before necropsy. Normalised data of the lung and muscle proteomes can be found in Supplementary Tab. 2-3.

The mass spectrometry proteomics data have been deposited to the ProteomeXchange Consortium via the PRIDE partner repository with the dataset identifier PXD070379.

### Metabolomics and Lipidomics extraction and identification

To perform untargeted metabolomics and lipidomics, ∼50 mg of frozen lung and muscle tissue were supplied to the Metabolomics Core Facility of the European Molecular Biology Laboratory, Heidelberg, Germany.

Sample extraction was initiated by adjusting tissue concentration homogenously across samples to 75 mg/mL via addition of 80% methanol, including internal standards (final conc.: 0.5% and 1.0%; for metabolomic profiling of small, polar molecules: MSK-A2-1.2, Cambridge Isotope Laboratories, USA; for lipidomics: EquiSPLASH, Avanti Polar Lipids, USA). After homogenisation on dry ice with a bead beater (FastPrep-24; MP Biomedicals, CA, USA) at 6.0 m/s (5 x 30 s, 5 min pause) using 1.0 mm zirconia beads (Biospec Products, OK, USA), samples were incubated at -20°C for 20 min. Subsequently, samples were vortexed and 187.5 µL per sample were transferred to a fresh 1.5 mL tube. For biphasic extraction of lipids and polar metabolites, 500 µL methyl tert-butyl ether were added. The monophasic mixture was vortexed for 60 s and incubated at -20°C for 20 min. For phase separation, 87.5 µL water were added, followed by another vortex and incubation step (see previous conditions). The biphasic solvent system was then centrifuged for 10 min at 15,000g and 4°C.

For lipidomics analysis, 400 µL of the upper organic phase were transferred, dried under a nitrogen stream and reconstituted in 200 µL isopropanol:methanol (50:50). For metabolomics analysis, 250 µL of the lower aqueous phase were transferred, dried under a nitrogen stream, and reconstituted in 100 µL ACN/methanol/H_2_O (2:2:1). The final samples were vortexed for 10 min, centrifuged (see previous conditions) and the supernatants were transferred to analytical glass vials for LC-MS/MS analysis.

The LC-MS/MS analysis was performed on a Vanquish Horizon UHPLC system, coupled to an Orbitrap Exploris 240 high-resolution mass spectrometer (Thermo Scientific, MA, USA) in negative and positive electrospray ionisation mode.

#### Untargeted Lipidomics

Chromatographic separation was carried out on an ACQUITY Premier CSH C18 column (2.1 mm x 100 mm, 1.7 µm, Waters, USA) at a flow rate of 0.3 mL/min. The mobile phase consisted of H_2_O:ACN (40:60; mobile phase A) and isopropanol:ACN (9:1; mobile phase B), which were modified with a total buffer concentration of 10 mM ammonium acetate and 0.1 % acetic acid. The following gradient (23 min total run time, including re-equilibration) was applied (min/%B): 0/15, 2.5/30, 3.2/48, 15/82, 17.5/99, 19.5/99, 20/15, 23/15. Column temperature was maintained at 65°C, the autosampler was set to 4°C and sample injection volume was 2 µL. Analytes were recorded via a full scan with a mass-resolving power of 12,0000 over a mass range from 200 – 1700 m/z (scan time: 100 ms, RF lens: 70%) in polarity-switching mode. To obtain MS/MS fragment spectra, data-dependant acquisition was carried out (resolving power: 15,000; scan time: 54 ms; stepped collision energies [%]: 25/35/50; cycle time: 600 ms). Ion source parameters were set to the following values: spray voltage: 3,250 V/ 3,000 V, sheath gas: 45 psi, auxiliary gas: 15 psi, sweep gas: 0 psi, ion transfer tube temperature: 300°C, vaporizer temperature: 275°C.

#### Untargeted Metabolomics (positive ionisation mode)

Chromatographic separation was carried out on an Atlantis Premier BEH Z-HILIC column (2.1 mm x 100 mm, 1.7 µm; Waters, USA) at a flow rate of 0.25 mL/min. The mobile phase consisted of H_2_O:ACN (9:1; mobile phase A) and ACN:H_2_O (9:1; mobile phase B), which were modified with a total buffer concentration of 10 mM ammonium formate. The aqueous portion of each mobile phase was pH-adjusted to pH 3.0 via addition of FA. The following gradient (20 min total run time, including re-equilibration) was applied (time [min]/%B): 0/95, 2/95, 14.5/60, 16/60, 16.5/95, 20/95. Column temperature was maintained at 40°C, the autosampler was set to 4°C and sample injection volume was 4 µL. Analytes were recorded via a full scan with a mass resolving power of 120,000 over a mass range from 60 – 900 m/z (scan time: 100 ms, RF lens: 70%). To obtain MS/MS fragment spectra, data-dependant acquisition was carried out (resolving power: 15,000; scan time: 22 ms; stepped collision energies [%]: 30/50/70; cycle time: 900 ms). Ion source parameters were set to the following values: spray voltage: 4,100 V, sheath gas: 30 psi, auxiliary gas: 5 psi, sweep gas: 0 psi, ion transfer tube temperature: 350°C, vaporizer temperature: 300°C.

#### Statistical Analysis

Lipidomics and metabolomics measurements were performed in separate analytical sequences, with experimental samples randomised within each sequence. Pooled QC samples were prepared by mixing equal aliquots from each processed sample. Multiple QCs were injected at the beginning of the analysis in order to equilibrate the analytical system. A QC sample was analysed after every 5^th^ experimental sample to monitor instrument performance throughout the sequence. For determination of background signals and subsequent background subtraction, an additional processed blank sample was recorded. Data was processed using MS DIAL 4.9.221218 (Tsugawa et al., 2015) and raw peak intensity data was normalised via total ion count of all detected features (Drotleff & Lacmmerhofer, 2019). Level 1 feature identification was based on the MS-DIAL LipidBlast V68 library (lipidomics) and an in-house library for metabolomics (EMBL-MCF 2.0; Dekina et al., 2024) using accurate mass, isotope pattern, MS/MS fragmentation, and retention time information and a minimum matching score of 80%. To remove misannotations and to enhance confidence in lipid identification, intra-class elution patterns of lipid species were checked for consistency by relying on the expected chromatographic behaviour on reversed phase columns within homologous lipid series, considering carbon chain length and degree of saturation as the main factors.

After data curation, features with a CV >30% QC samples were excluded from the results. For univariate statistical analysis, data were log-transformed and unpaired t-tests were conducted between experimental groups of interest. To control for type I errors resulting from multiple testing, p-values were adjusted using the SGOF method (Carvajal-Rodríguez et al., 2009), and only features with an adjusted p-value < 0.05 were considered significant.

Raw data of the metabolomics and lipidomics analyses are available at the NIH Common Fund’s National Metabolomics Data Repository website, the Metabolomics Workbench, under the study ID ST004261. Normalised data of the lung and muscle metabolomes and lipidomes are provided in Supplementary Tab. 4-9.

### Differential and Pathway Analysis and Visualisation

Differentially abundant proteins, metabolites and lipids were defined by two different threshold cut-offs.

To identify the most significant differentially abundant features in non-healthy compared to healthy harbour porpoises, only proteins, metabolites and lipids were considered, if adjusted p-value (pval_adj_) ≤ 0.05 and if logarithmic fold change (FC_log2_) ≥ 0.58 for more abundant or ≤ -0.58 for less abundant.

For Gene Ontology (GO) term and pathway analysis, and for lipid subclass analysis, proteins, metabolites and lipids with an unadjusted p-value ≤ 0.05 and FC_log2_ ≥ 0.58 or ≤ -0.58nwere included to get an overview of the biological context between the two conditions.

If multiple detections of the same lipid were identified due to the different adducts of the detection in positive and negative mode, only lipids with the adduct “[M+H]^+^” or “[M+H]^-^“ were kept. If a lipid was detected with both of these adducts, only the “[M+H]^+^”-associated lipid was kept.

The proteins were analysed for statistical overrepresentation of annotated GO-Slim terms in “Biological Processes” with the PANTHER classification system (reference: *Homo sapiens*; version 19.0, Thomas et al., 2022). Fisher’s exact test was used for FDR correction. Only GO-Slim terms with the most specific child term were selected instead of broader umbrella terms. For the overrepresentation analysis of the metabolites, MetaboAnalyst was used and compared against the RaMP database (version 6.0, Pang et al., 2024). Only metabolic reference pathway sets were used that contained at least two metabolite entries and statistically significant results were FDR-corrected. Significantly dysregulated lipids were analysed on a subclass level with LipidOne (reference: *H. sapiens*; version 2.3, Alabed et al., 2024). Statistical significances of differential lipid subclasses between the two conditions, non-healthy and healthy, were determined with a t-test and corrected for FDR.

### Omics Integration

For omics integration, all individuals (n = 12) with corresponding proteomes, metabolomes and lipidomes of the lung and the muscle were included. Omics integration was performed with “mixOmics” R package (Rohart et al., 2017) and the vignette was followed as advised for supervised n-integration with “DIABLO” (Data Integration Analysis for Biomarker Discovery using Latent Variable Approaches for Omics Studies). Normalised features of the proteomes, metabolomes and lipidomes were included, if p-value ≤ 0.05 and not further filtered. For multiple detections of the same lipid with different adducts, the same filtering procedure was followed as stated previously. Sparse Partial Least Square-Discriminant Analysis (sPLS-DA) was performed to classify the samples and identify the most crucial features for discrimination. Model tuning steps were set to include centroids as distance metric and the “Leave One Out” (LOO)-method with five folds and 50 repeats for cross-validation. Significant features and signatures were cut-off at a correlation coefficient of r > 0.7 which indicates strong correlation.

## RESULTS

### Identification and sensitivity of omics on post-mortem harbour porpoise samples

Using a LC-MS/MS approach, we identified a total of 2560 proteins, 109 metabolites and 1093 lipids in the lung, and 1526 proteins, 110 metabolites and 1101 lipids in the muscle across non-healthy and healthy harbour porpoises. Of the identified features, only 33 proteins, two metabolites and 106 lipids in the lung and 21 proteins, 11 metabolites and 60 lipids in the corresponding muscles showed significant dysregulation in non-healthy compared to healthy individuals (pval_adj_ ≤ 0.05, FC_log2_ ≥ 0.58 or ≤ -0.58; Fig. 1; Tab. 2).

**Figure 1.**
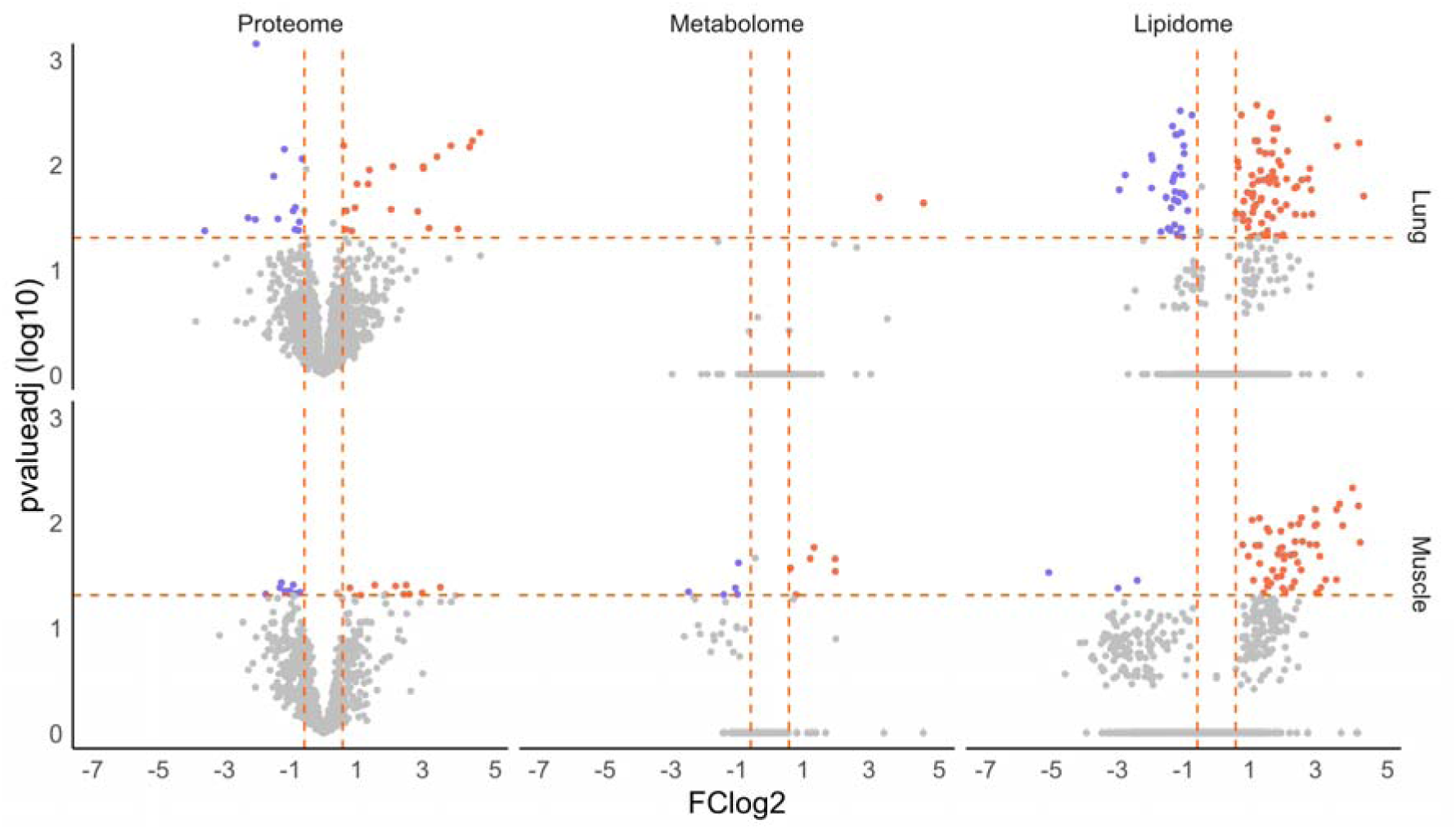
Differential proteins, metabolites and lipids in the lung and muscle of non-healthy, wild harbour porpoises. The x-axis indicated the log2-transformed fold change (FC_log2_) and the y-axis the log10-transformed adjusted p-value. The dashed lines mark the significance cut-offs of the adjusted p-value (pval_adj_ ≤ 0.05) and FC_log2_ (higher abundance: FC_log2_ ≥ 0.58, lower abundance: FC_log2_ ≤ -0.58). Features are coloured according to their significant dysregulation in non-healthy harbour porpoises compared to healthy individuals: purple for less abundant features and orange for more abundant. A total of 111 not significant metabolites and lipids had FC_log2_ values outside of the here depicted range but were omitted from the graph to enhance visualisation (pval_adj_ ≥ 0.05; lowest FC_log2_ = -13, highest FC_log2_ = 18).

**Table 2.** Overview of applied cut-offs and features included in the analyses. Cut-offs of the normalised features for the differential analysis, the pathway and GO analysis and the omics integration are defined. With the applied cutoffs, the number of proteins, metabolites and lipids in the lung and muscle that were considered for each analysis is stated. For the differential analysis and the pathway and GO analysis, the regulation in nonhealthy harbour porpoises compared to healthy animals is specified.

|  |  | Dysregulated Features |  | Pathway and GO Analysis |  | Omics Integration |
| --- | --- | --- | --- | --- | --- | --- |
|  | Cutoffs | FC<sub>log2</sub> ≥ 0.58 / ≤ -0.58; pval<sub>adj</sub> ≤ 0.05 |  | FC<sub>log2</sub> ≥ 0.58 / ≤ -0.58; p-value ≤ 0.05 |  | p-value ≤ 0.05 |
|  |  | up | down | up | down |  |
| Lung | Proteins | 20 | 13 | 114 | 104 | 2037 |
|  | Metabolites | 2 | 0 | 9 | 2 | 112 |
|  | Lipids | 72 | 34 | 116 | 50 | 1082 |
| Muscle | Proteins | 10 | 11 | 70 | 116 | 1135 |
|  | Metabolites | 6 | 5 | 7 | 17 | 112 |
|  | Lipids | 57 | 3 | 105 | 16 | 1091 |

### MultiOmics analysis of the lung of non-healthy harbour porpoises

#### Proteins showed enhanced metabolic and immune processes and reduced ECM integrity

In the lung, 20 proteins were significantly enhanced, whereas 13 proteins showed significantly lower abundance in the non-healthy compared to healthy harbour porpoises (Fig. 1; Tab. 2). We identified the top up- and downregulated GO-Slim Biological Processes. Processes related to metabolism, such as purine metabolism and monocarboxylic acid metabolism, were highly elevated in non-healthy harbour porpoises (Fig. 2A, Tab. 3). Various highly abundant proteins possessed a role in oxidative stress responses, including in glutathione metabolism (GPX1, PRDX6, GPX3, ERO1A, GSTP1, SOD2, TXNRD1; Fig. 2A, Tab. 3; Supplementary Tab. 2), and innate immune responses (CTSB, DPP7, PDXK, PRDX6, C3, FLT, TCIRG1, C8A, PIN1, C4BPA, C7, PSAP, CREG1, GSTP1, PSMA4, SFTPD, MAN2B1, C6, S100A8, EPX, LAMP1, FABP5, S100A12, PNP, CTSD; Fig. 2A, Tab. 3; Supplementary Tab. 2). The most significant reduction was found for NFκB signalling transduction, protein stability and modification processes (Fig. 2A). Other significantly reduced processes and decreased proteins were associated with extracellular matrix and transcription, such as splicing or chromatin assembly (Fig. 2A, Tab. 3).

**Figure 2.**
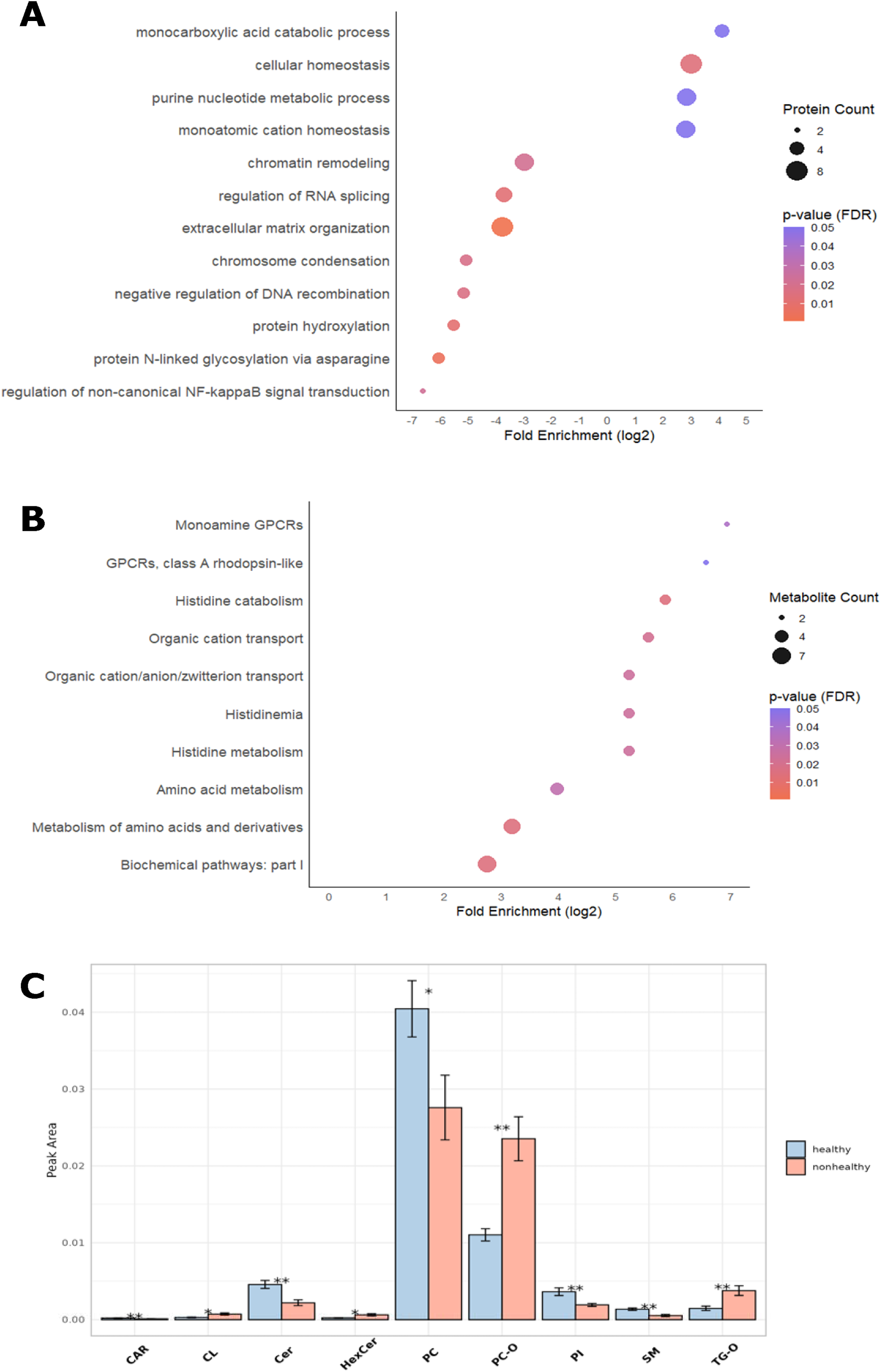
GO terms (proteins), pathways (metabolites) and lipid subclasses in the lung of non-healthy harbour porpoises compared to healthy individuals. Proteins, metabolites and lipids with a cut-off of p-value ≤ 0.05, higher abundance: FC_log2_ ≥ 0.58, lower abundance: FC_log2_ ≤ -0.58 were considered. A Proteins were analysed for overrepresented GO-Slim “Biological process” terms using PANTHER against the reference dataset of the human. The term “Biochemical pathway: part I” encompasses all primarily metabolic pathways in mammals. B Metabolites were analysed with MetaboAnalyst for overrepresented metabolic pathways against the RaMP database. For A and B, the number of involved proteins or metabolites found differentially abundant in non-healthy harbour porpoises is depicted by increasing size of the dot. Significance of the respective pathways and processes is indicated by a colour scale. Fold enrichment was log2-transformed. C Lipid subclass composition of significantly altered lipids between the conditions was analysed with LipidOne. Not significant subclasses were excluded. The peak area is a measure for lipid abundance. Significant differences between conditions were calculated with an ANOVA or t-test and are indicated by asterisks (*: < 0.05, ** < 0.01). CAR: carnitine, CL: cardiolipin, CER: ceramide, HexCer: hexosylceramide, PC: phosphatidylcholine, PC O-: ether-phosphatidylcholine, PI: phosphatidylinositol, SM: sphingomyelin, TG O-: ether-triacylglycerol.

**Table 3.**
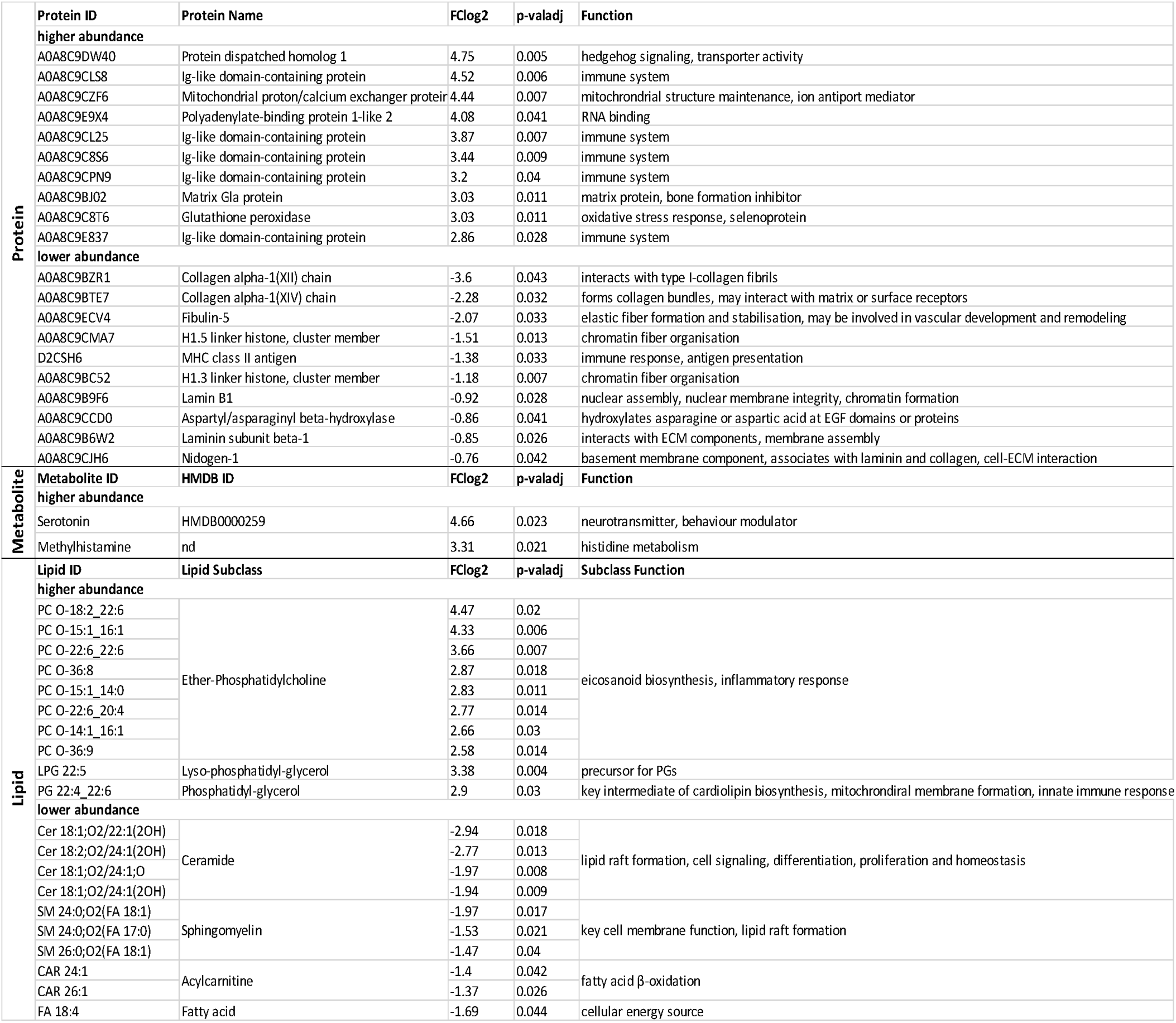
Top 10 significantly dysregulated proteins, metabolites and lipids in the lung of non-healthy harbour porpoises. Proteins, metabolites and lipids were considered highly significantly dysregulated if adjusted pvalue ≤ 0.05 and logarithmic fold change (FC_log2_) ≥ 0.58 for more abundant and ≤ -0.58 for less abundant features in non-healthy porpoises in comparison to healthy porpoises. Protein IDs, names and function were inferred from the UniProt protein database for the vaquita (Phocoena sinus). Metabolites and lipids were identified with MS-Dial against a metabolite and lipid database. For metabolites, the identification number of the Human Metabolite database (HMDB) is given, which was used to define the function. The function of methylhistamine was not exactly determinable and was inferred from 1-methylhistamine (HMDB0000898). For the lipids, only the function of the lipid subclass of significantly dysregulated lipids could be given. To describe the function, the lipid subclass descriptions of lipotype.com were used. The statistical significance was calculated with a student’s t-test for proteins and with the sequential goodness of fit test for metabolites and lipids.

#### Metabolites showed elevated histamine and histidine metabolism

In lungs of non-healthy harbour porpoises, metabolic pathways related to amino acid metabolism, such as histidine, and transport processes (e.g., monoamine GPCRs, organic cation transport) were elevated (Fig. 2B). Only serotonin and methylhistamine were significantly elevated in non-healthy harbour porpoises compared to the healthy control (Tab. 3).

#### Lipids showed high abundance of ether-linked lipids and reduced lipids integral to membrane and lipid raft formation

The analysis of differential lipid subclasses with LipidOne indicated a significantly altered composition for nine of 19 dysregulated subclasses between non-healthy and healthy porpoises (Fig. 2C). These included ether-linked lipid subclasses, such as ether-phosphatidylcholine (PC O-), that showed high elevation in non-healthy individuals (Fig. 2C, Tab. 3). Cardiolipins along with intermediate lipids, were also found significantly elevated in non-healthy porpoises (LPG 22:5, PG 22:4_22:6; Fig. 2C, Tab. 3). Significant reduction was found for lipids involved mainly in membrane integrity and fluidity (PC, ceramide, sphingomyelin, phosphatidylinositol) and fatty acid metabolism (carnitine, free fatty acid; Fig. 2C, Tab. 3).

### MultiOmics analysis of the muscle of non-healthy harbour porpoises

#### Proteins showed elevated muscle structure pathways and decreased **β**-oxidation

Significantly elevated proteins were mainly involved in muscle function and structure (Fig. 3A; Tab. 4). Multiple PDLIM family proteins with a muscular structure function (PDLIM7, LDB3, PDLIM5) and transport-related proteins were significantly more abundant in muscles of non-healthy harbour porpoises (Tab. 4). The highest upregulation in the muscle was found for the same protein in the lung, protein dispatched homolog 1 (Tab. 3, 4). Various biological processes were significantly reduced, including metabolism, translation and mitochondrial structure (Fig. 3A). Most downregulated terms were associated with lipid metabolism, specifically β-oxidation (STOML2, FABP3, ETFDH) and including both subunits of fatty acid metabolism key mediator, trifunctional enzyme (HADHA, HADHB; Fig. 3A, Tab. 4). Further reduced processes included protein folding and translation (Fig. 3A, Tab. 4) and the most significantly reduced proteins were chaperone complex proteins that assist in correct protein assembly (CCT5, CCT2; Tab. 4).

**Figure 3.**
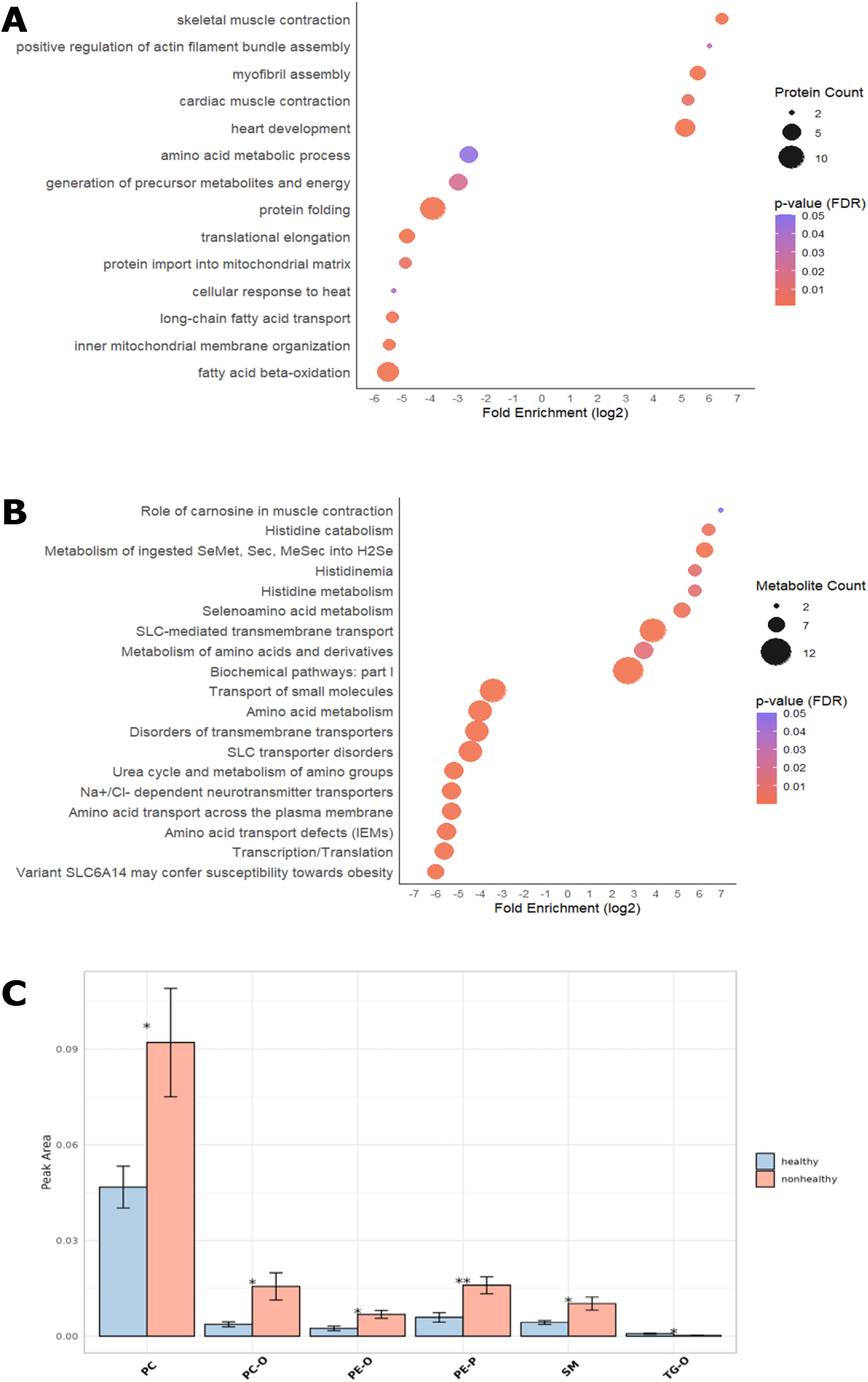
GO terms (proteins), pathways (metabolites) and lipid subclasses in the muscle of non-healthy harbour porpoises compared to healthy individuals. Proteins, metabolites and lipids with a cut-off of p-value ≤ 0.05, higher abundance: FC_log2_ ≥ 0.58, lower abundance: FC_log2_ ≤ -0.58 were considered. A Proteins were analysed for overrepresented GO-Slim “Biological process” terms using PANTHER against the reference dataset of the human. B Metabolites were analysed with MetaboAnalyst for overrepresented metabolic pathways against the RaMP database. For A and B, the number of involved proteins or metabolites found differentially abundant in non-healthy harbour porpoises is depicted by increasing size of the dot. Significance of the respective pathways and processes is indicated by a colour scale. Fold enrichment was log2-transformed. C Lipid subclass composition of significantly altered lipids between the conditions was analysed with LipidOne. Not significant subclasses were excluded. The peak area is a measure for lipid abundance. Significant differences between conditions were calculated with an ANOVA or t-test and are indicated by asterisks (*: < 0.05, ** < 0.01). PC: phosphatidylcholine, PC O-: ether-phosphatidylcholine, PE O-: oxidised ether-phosphatidylethanolamine, PE P-: ether-phosphatidylethanolamine, SM: sphingomyelin, TG O-: ether-

**Table 4.**
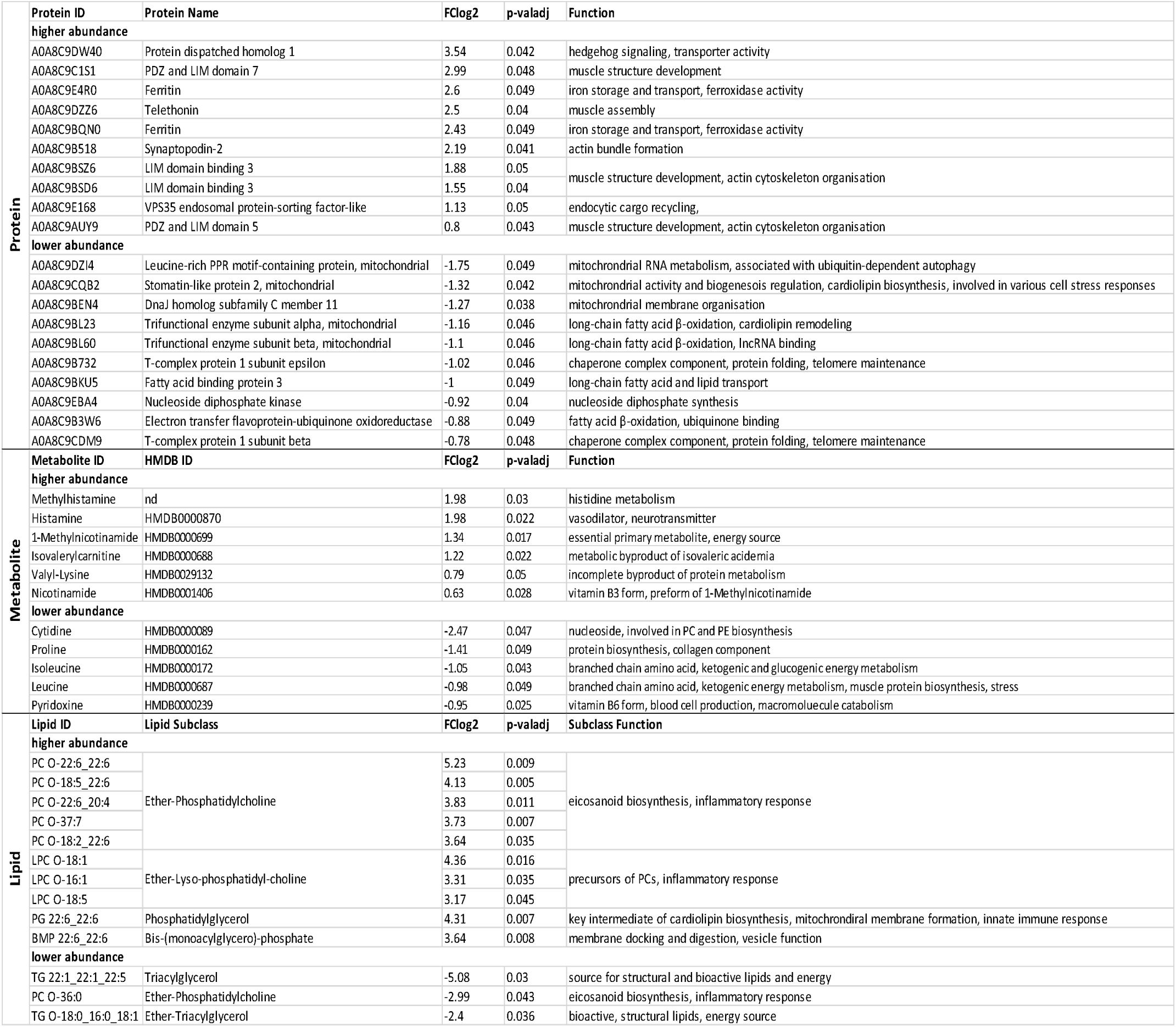
Top 10 significantly dysregulated proteins, metabolites and lipids in the muscle of nonhealthy harbour porpoises. Proteins, metabolites and lipids were considered highly significantly dysregulated if adjusted p-value ≤ 0.05 and logarithmic fold change (FC_log2_) ≥ 0.58 for more abundant and ≤ -0.58 for less abundant features in non-healthy porpoises in comparison to healthy porpoises. Protein IDs, names and function were inferred from the UniProt protein database for the vaquita (Phocoena sinus). Metabolites and lipids were identified with MS-Dial against a metabolite and lipid database. For metabolites, the identification number of the Human Metabolite database (HMDB) is given, which was used to define the function. The function of methylhistamine was not exactly determinable and was inferred from 1-methylhistamine (HMDB0000898). For the lipids, only the function of the lipid subclass of significantly dysregulated lipids could be given. To describe the function, the lipid subclass descriptions of lipotype.com were used. The statistical significance was calculated with a student’s t-test for proteins and with the sequential goodness of fit test for metabolites and lipids.

#### Metabolites showed dysregulation of incomplete protein metabolism and transport processes

Muscles of non-healthy harbour porpoises were mostly enriched for histidine and histamine metabolism and selen-associated metabolic pathways (Fig. 3B, Tab. 4). Similar to the proteome, dysregulated metabolites in the muscles of non-healthy harbour porpoises showed a reduced enrichment for amino acid metabolism, transcriptional and translational processes. Most terms that showed a significant reduction were associated with transport activities (e.g., amino acid transport defects, Na^+^/Cl^-^-dependent neurotransmitter transporters). In concordance, instable and intermediate protein metabolism metabolites, valyl-lysine and isovalerylcarnitine, were also highly enhanced (Tab. 4). Moreover, isoleucine, leucine and cytidine, metabolites related to fatty acid metabolism, exhibited lower abundance (Tab. 4).

#### Lipids showed phospholipids and sphingomyelins as highly abundant

Significantly differentially abundant lipids were classified into a total of 20 identified lipid subclasses. Of these, six showed a significant difference between conditions (Fig. 3C). Except for ether-triacylglycerols (TC O-), all lipid subclasses were significantly more abundant in non-healthy harbour porpoises. These included main membrane lipids, such as phosphatidylcholines and sphingomyelins. Plasmalogens (PC O-, PE O-, PE P-) showed a high increase in the muscles of non-healthy individuals compared to healthy porpoises (Fig. 3C, Tab. 4).

#### Dysregulated cross-tissue features in the lung and muscle of non-healthy harbour porpoises

We analysed features for similar dysregulation in both, the lung and the muscle (p-value ≤ 0.05; FC_log2_ ≥ 0.58 and ≤ -0.58). Figure 4 depicts Venn diagrams that illustrate cross-tissue dysregulated features for each omics layer. Comparing dysregulated proteins from the lung with those of the muscle, 13 proteins showed upregulation in both and five were downregulated. Interestingly, five of the upregulated proteins across both tissues play an important role in antioxidant defence and detoxification (PRDX6, GLO1, GSTP1, ferritin; Fig. 4A). Moreover, five immunoglobulins showed an increase in abundance in both tissues. Notably, four of these immunoglobulins were also among the most elevated proteins in the lung, while both ferritins were under the top 10 upregulated proteins in the muscle (Tab. 3, 4). CTSD is involved in protein breakdown, HINT1 is a hydrolase associated with purine nucleotides. The five concordantly decreased proteins are involved in nuclear processes and cell cycle (SFPQ, CENPV) and membrane and ECM stability (DNAJC11, HSPG2, SERPINH1). Proteins that exhibited inconsistent cross-tissue regulation and were upregulated in the lung but downregulated in the muscle were involved in metabolite detoxification (ECHDC1), structure maintenance (LETM1, DSTN) and β-oxidation (ECI2).

**Figure 4.**
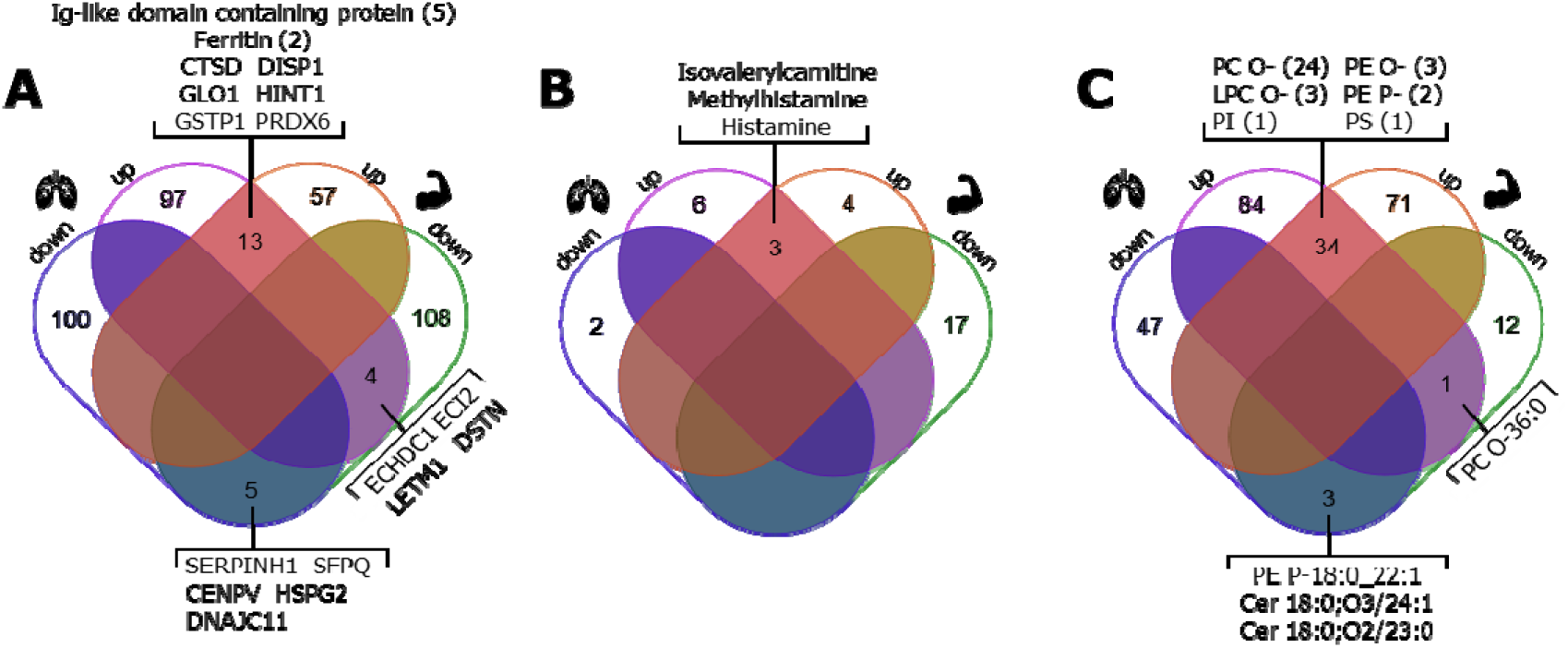
Venn diagrams of similarly dysregulated proteins, metabolites and lipids between the lung and muscle. Features were considered if p-value < 0.05 and FC_log2_ > 0.58 (more abundant) or < -0.58 (less abundant) in the tissues of non-healthy compared to healthy harbour porpoises. On the left side of the Venn diagrams, the upregulated (purple) and downregulated (blue) features are shown, while on the right side, upregulated (orange) and downregulated (green) features of the muscle are highlighted. A depicts similarly dysregulated proteins in the lung and muscle. Of the 22 identified proteins, 13 were upregulated in both tissues and five proteins were found to be decreased in the lung and muscle. Four proteins showed dissimilar regulation and were upregulated in the lung, but downregulated in the muscle. The numbers in brackets behind ferritin and Ig-like domain containing protein indicate the number of unique proteins that were identified with this name but could not be further specified. B shows overlapping metabolites in both tissues. Only three cross-tissue metabolites were identified, all of which were upregulated in both tissues. C shows the Venn diagram for the lipids of the lung and muscle. Due to the large number, cross-tissue upregulated lipids were summarised into lipid subclasses. Of a total of 38 identified lipids, 34 were elevated in both tissues and encompassed six subclasses. Three lipids were found decreased in both tissues and one lipid exhibited dissimilar regulation. The numbers in brackets represent the number of identified lipids of this subclass, respectively. Cer: ceramide, PC O-: ether-phosphatidylcholine, PE O-: oxidised ether-phosphatidylethanolamine, PE P-: ether-phosphatidylethanolamine, PI: phosphatidylinositol, PS: phosphatidylserine.

Histamine, methylhistamine and isovalerylcarnitine were identified to be highly elevated metabolites in the lung and in the muscle of non-healthy harbour porpoises and were also among the significantly upregulated metabolites in the muscle (Fig. 4B; Tab. 4).

A total of 38 lipids were found similarly dysregulated in both tissues in non-healthy harbour porpoises, of which the majority was highly elevated (Fig. 4C). The 34 elevated lipids were summarised to subclasses, which were mainly ether-linked lipid classes. Most identified lipids belonged to ether-phosphatidylcholines (24 lipids) and to a lesser extent to other plasmalogens (5 lipids) or subclasses (LPC O-, PI, PS: phosphatidylserine). Two ceramides and one plasmalogen were less abundant in both tissues, while one ether-phosphatidylcholine exhibited nonconcurrent regulation across both tissues. plasmalogen were less abundant in both tissues, while one ether-phosphatidylcholine exhibited nonconcurrent regulation across both tissues.

#### Integrated MultiOmics analysis and signature in the lung and muscle

By integrating normalised abundance data of all omics layers for each tissue with mixOmics “DIABLO”, we defined key features for a signature that best explains the molecular differentiation between non-healthy and healthy harbour porpoises. Since our sample size consisted of six individuals in each group, we restricted the number of contributing features to six in the top two components for each omics set. The discrimination between non-healthy harbour porpoises and healthy harbour porpoises was assessed with a sPLS-DA (Fig. 5A, 6A). Clustering of the conditions was more discrete in the lung than in the muscle, especially for the metabolite and lipid blocks (Fig. 5A, 6A). One non-healthy harbour porpoise, Pph_nh6, and one healthy harbour porpoise, Pph_h3, clustered closely in the lung analysis. Similarly, for the metabolites and lipids in the muscle, two non-healthy (Pph_nh3, Pph_nh6) and two healthy (Pph_h3, Pph_h6) samples showed slight clustering. Next, we assessed the correlation between the omics layers (Fig. 5B, 6B). The correlation between the omics sets of the lung was high, with correlation coefficients around 0.9 indicating an appropriate design matrix that distinguishes between the health conditions (Fig. 5B). Similar to the results of the sPLS-DA, the correlation of the muscle was lower compared to the lung, but still had high correlation coefficients of > 0.8 (Fig. 6B). The abundance profiles for all individuals further illustrate the separation of the conditions based on the five most contributing proteins, metabolites and lipids on the first component (Fig. 5C, 6C; Tab. 5). While there was a slight inter-individual variation retained for various features, mostly for metabolites including serotonin, isovalerylcarnitine and taurine in the lung, and proteins, such as A0A8C9CLS8 and A0A8C9CL25 (both encoding for Ig-like domain-containing proteins) in the muscle, the health conditions exhibited clear clustering (Fig. 5C, 6C). Again, only two samples, Pph_nh6 and Pph_h3, clustered closely together for both tissues (Fig. 5C, 6C). Lastly, we analysed the connection between the six most contributing features across the omics layers (Fig. 5D, 6D, Tab. 5, Supplementary Tab. 10). For this, we set a correlation cut-off at *r* > 0.7. Proteins, metabolites and lipids showed strong, both positive and negative, interconnected links in the lung and muscle (Fig. 5D, 6D). In the lung, A0A8C9CL25 (encoding an Ig-like domain-containing protein), taurine and two ceramides (Cer 18:1;O2/22:0;O, Cer 18:0;O2/22:0) were mostly associated with negative correlations, while SLC25A4 was only associated with positive correlations to TG O-16:0_16:0_20:4 and O-phosphoethanolamine (Fig. 5D, Tab. 5). In the muscle, anserine showed no significant correlation with any of the most contributing features. A high, positive correlation was identified for SPEG and BMP 18:1_22:6, which were both negatively correlated with pipecolic acid (Tab. 5). Notably, many metabolites (e.g., pipecolic acid, leucine, isoleucine, proline) were only negatively correlated with other contributing features (Fig. 6D).

**Figure 5.**
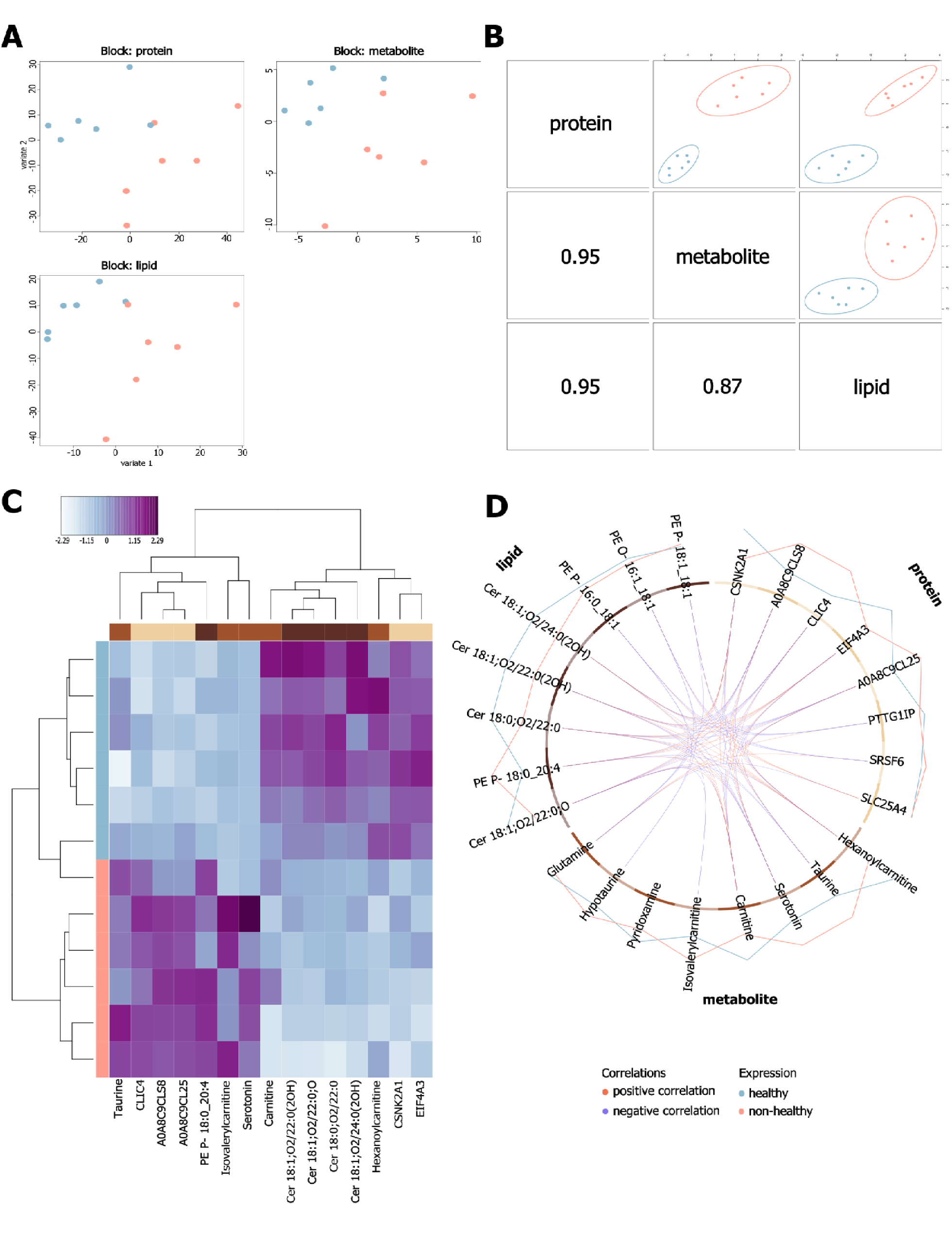
Integrative MultiOmics of the lung using DIABLO. Across all graphics, apricot represents non-healthy individuals, light blue depicts healthy animals. A shows the sPLS-DA plot of each omics layer used for omics integration, projected on both components (variant 1, variant 2). The sPLS-DA illustrates the discrimination calculated on the most contributing features between the health conditions. One non-healthy and one healthy animal share similar abundance profiles for the proteins and lipids. B depicts the correlations between abundance matrices of the different omics for component 1 based on the most contributing features. The plots illustrate the maximised discrimination power to distinguish between health conditions. The left-bound numbers represent the correlation coefficients of the respective pair of omics set (protein-metabolite: 0.95; metabolite-lipid: 0.87; protein-lipid: 0.95). A correlation coefficient over 0.8 indicates high correlation between omics sets. C Heatmap of the most discriminative features in the lung of non-healthy and healthy harbour porpoises. Individual samples are coloured vertically on the left and names of the contributing features are given at the bottom. Features types are colour-coded: proteins are beige, metabolites are brown and lipids are dark brown. Differential abundance of the features is also indicated by a colour gradient from white (low) to purple (high). D shows a circos plot of the correlations of the six features contributing the most to the discrimination between health conditions. Only correlations with a greater correlation coefficient of 0.7 are shown. The colour blocks each represent a different omics set (proteins: beige, metabolites: brown, lipids: dark brown). The inner lines visualise the correlations between the features, with dark orange purple indicating positive and indicating negative correlations. The outer encompassing lines represent the abundance levels of the features in the lungs of non-healthy (orange line) and healthy (blue line) harbour porpoises.

**Figure 6.**
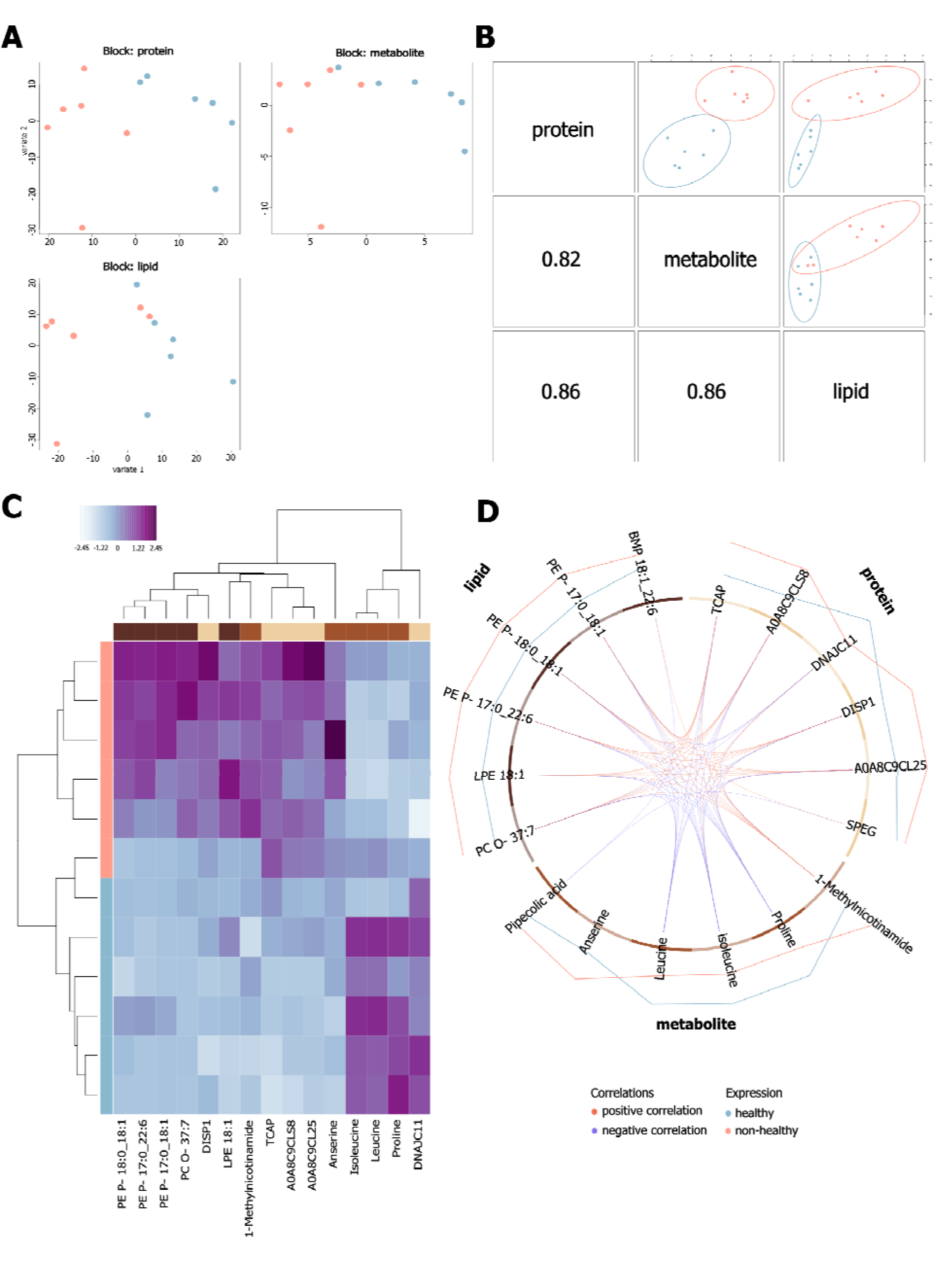
Integrative MultiOmics of the muscle using DIABLO. Across all graphics, orange represents non-healthy individuals, blue depicts healthy animals. A shows the sPLS-DA plot of each omics layer used for omics integration, projected on both components (variant 1, variant 2). Individual identification numbers of the harbour porpoises are given. The sPLS-DA illustrates the discrimination calculated on the most contributing features between the health conditions. Across the metabolites and lipids, two non-healthy and two healthy animals share similar abundance profiles. B depicts the correlations between abundance matrices of the different omics for component 1 based on the most contributing features. The plots illustrate the maximised discrimination power to distinguish between health conditions. The left-bound numbers represent the correlation coefficients of the respective pair of omics set, with values of > 0.8 indicating high correlation (protein-metabolite: 0.82; metabolite-lipid: 0.86; protein-lipid: 0.86). C Heatmap of the most discriminative features in the muscle of non-healthy and healthy harbour porpoises. Individual samples are coloured (orange: non-healthy, blue: healthy) and names of the contributing features are given. Features types are colour-coded: proteins are beige, metabolites are brown and lipids are dark brown. Differential abundance of the features is also indicated by a colour gradient from white (low) to violet (high). D shows a circos plot of the correlations of the six features contributing the most to the discrimination between health conditions. Only correlations with a correlation coefficient of > 0.7 are shown. The blocks each represent a different omics set (proteins: beige, metabolites: brown, lipids: dark yellow). The inner lines visualise the correlations between the features, with dark orange dark orange indicating positive and purple indicating negative correlations. The outer encompassing lines represent the abundance levels of the features in the muscle of non-healthy (orange line) and healthy (blue line) harbour porpoises.

**Table 5.**
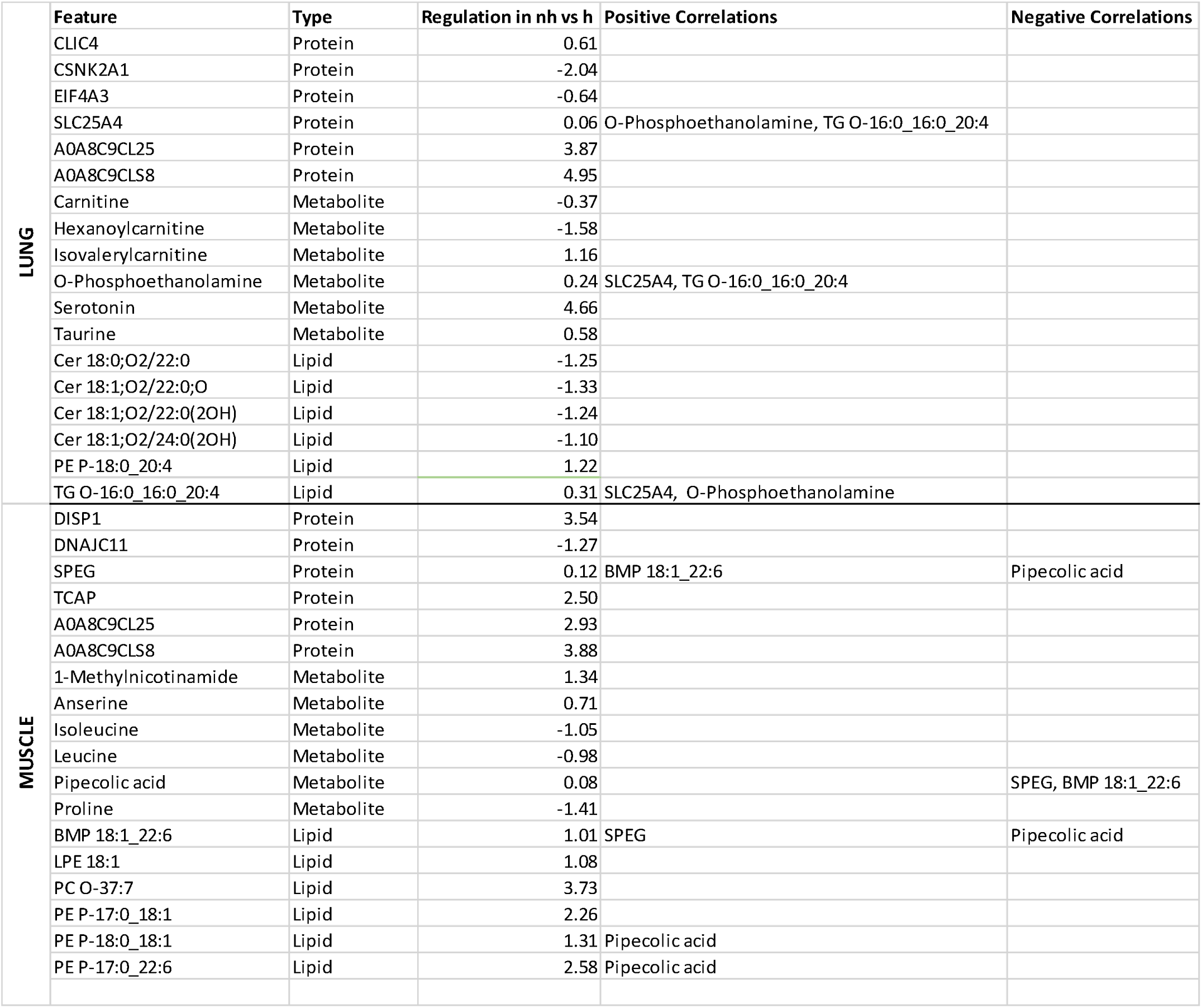
Correlation signature of the lung and muscle in non-healthy harbour porpoises. Correlations between the features across the omics data were identified with mixOmics DIABLO tool which can be utilised to describe a specific marker fingerprint of a condition or disease. The regulation found in the respective omics data sets is given as logarithmic fold change (FC_log2_). Positive and negative correlations of the features, as identified with DIABLO and visualised in Fig. 5D and 6D, is noted as well. Within the identified correlated signatures, one smaller cluster was identified in for each tissue. In the lung, this signature encompassed SLC25A4, O-phosphoethanolamine and TG O-16:0_16:0_20:4. The muscle signature combined positively correlated features SPEG and BMP 18:1_22:6 which were negatively correlated with pipecolic acid. Pipecolic acid was positively correlated with PE P-18:0_18:1 and PE P- 17:0_22:6. For a more detailed table with correlation coefficients, see Supplementary Tab. 11.

## DISCUSSION

### Necropsy material suitable for omics studies

In this study, proteins, metabolites and lipids of the lung and the muscle of free-ranging harbour porpoises, that were sampled during routine necropsies, were extracted and analysed using LC-MS/MS methods. With this omics approach, a total of 2560 proteins, 109 metabolites and 1251 lipids in the lung, and 1526 proteins, 110 metabolites and 1259 lipids in the muscle were identified. As of date, there is no comparable study of marine mammal lungs, as this is not an easily accessible and hard-to-work-with tissue. However, proteomic, metabolomic and lipidomic studies have been performed on breath exhale (blow) samples of trained bottlenose dolphins, though with less identified features (Aksenov et al., 2014; Bergfelt et al., 2018; Pasamontes et al., 2017). Many factors can influence omics analyses and yield, including extraction and detection protocols, distribution within the tissue and exact tissue sample localisation (Kershaw et al., 2018). Additionally, marine mammal reference libraries are often incomplete and still subject to ongoing studies (Mancia, 2018; Van Cise et al., 2024). This study therefore not only underlines the suitability of necropsy tissue material for omics analyses which can give timely insights into responses to environmental changes, but also adds valuable input as a reference for future studies.

### MultiOmics of the lung

In the lungs of non-healthy harbour porpoises, proteins and lipids involved in anti-inflammatory and antioxidative responses were highly abundant. This included glutathione-associated proteins (GPX1, GPX3, GSTP1) among other antioxidants, such as SOD2 and PRDX6, whose important roles in protection from hypoxia and oxidative insult in marine mammals are well described (Allen & Vázquez-Medina, 2019; Cantú-Medellín et al., 2011; Tian et al., 2019, 2021a, 2021b). Diseases represent an extraordinary stress, which often also causes metabolic alterations (Kivimäki et al., 2023). Here, we found purine metabolism-associated proteins elevated (HPRT1, PRPS1, ADSL, GMPR2). Similarly, enhanced purine metabolism was observed in stressed northern elephant seals (ACTH: Deyarmin et al., 2020; fasting: Soñanez-Organis et al., 2012) and in dolphins (del Castillo Velasco-Martínez et al., 2016). A higher utilisation of the purine salvage pathway has been proposed for marine mammals, which enables them to directly recycle hypoxanthine, a reactive oxygen species-generating product of purine metabolism, for ATP production (Soñanez-Organis et al., 2012). Likewise, energetically costly translational processes were reduced in lungs of non-healthy harbour porpoises, which is often one of the first reduced processes under stress or disease to prioritise energy resources (Shah et al., 2013).

High immunoglobulin concentrations have been observed in wild animals compared to marine mammals in human care, which is presumably due to lower pathogenic impact of managed animals (Fair et al., 2017). Immunoglobulins recognise and neutralise pathogens, toxins and other foreign particles (Schroeder & Cavacini, 2010) and were also identified in a proteomic study of blow from dolphins (Bergfelt et al., 2018). In line with this, we observed high abundance of immune system proteins, including highly abundant immunoglobulins, though many could not be precisely identified due to incomplete reference data. Additionally, inflammation-alleviating lipids (plasmalogens, Brosche & Platt, 1998) and metabolites were elevated in non-healthy harbour porpoises (serotonin, methylhistamine; Herr et al., 2017; Jutel et al., 2009). With the lung as a primary interface of the organism with the environment, thus with potential environmental insults, an activation of the immune system was expected in harbour porpoises diagnosed with lung diseases. However, contrary to the anti-inflammatory T helper cell 2 (Th2) response typically induced by pulmonary parasite infections, non-healthy harbour porpoises showed immune responses consistent with proinflammatory type 1 activation (Allen & Sutherland, 2014; Romagnani, 2006). Given that these individuals were diagnosed with severe nematode infestations and bronchopneumonia, this finding aligns with previous studies on harbour porpoises, reporting diminished Th2 responses in severe stages (Tsang et al., 2025), possibly reflecting an immune overcompensation or a compromised immune system.

The anatomy of the cetacean lung is specialised for life under water (Piscitelli et al., 2010, 2013; Reidenberg & Laitman, 2025). Their lungs and conducting airways are enforced with thickened alveolar walls and abundant collagenous elastic fibres (Piscitelli et al., 2010, 2013). While this phenotype is indicative of pulmonary fibrosis in the human (Crouch, 1990), in whales, this is an essential diving adaptation (Kooyman & Sinnett, 1979; Piscitelli et al., 2010, 2013). Collagens are building blocks of the extracellular matrix and play a role in wound healing (Stamenkovic, 2003). Here, we found underrepresented extracellular matrix processes and ten collagens decreased in the lungs of non-healthy harbour porpoises, indicating instability of the connective tissue which is necessary to facilitate gas exchange and lung collapse. Likewise, the lungs exhibited higher abundance of cardiolipins which have been associated with lung injury and impaired surfactant function in human cell culture models of pneumonia (Ray et al., 2010; Tyurina et al., 2010). Recent similar findings in dolphin lungs further support our observation and underscore the importance of accumulated cardiolipins in impacted lungs (Porras-Gómez et al., 2025). Lung surfactant not only facilitates gas exchange, but also clears foreign compounds and pathogens from the alveolar space (Whitsett et al., 2010). Moreover, surfactant proteins have been found to be under positive selection and accelerated evolution in marine mammals (Chikina et al., 2016). While surfactant composition and concentration can be indicative of respiratory diseases, it is also altered in drowning pathology (Lorente et al., 1990) which can be applied to by-caught marine mammals. However, we could not confirm a clear distinction in surfactant protein abundance (SFTPA, SFTPB, SFTPD) between by-caught and not by-caught porpoises.

The lipid profiling showed higher abundance of ether-phosphatidylcholines and ether-triacylglycerols compared to healthy individuals. It has been stated that the cetacean lung profile is composed mainly of phosphatidylcholines, phosphatidylethanolamines, phosphatidylserines, phosphatidylinositols and sphingomyelins, regardless of their diving abilities (Arregui et al., 2020). However, lungs of non-healthy harbour porpoises displayed significantly lower abundance of phosphatidylcholines, phosphatidylinositols and sphingomyelins compared to their healthy counterparts. This may point to an altered lipid composition and inflammatory state in the pathological lung (Mehta et al., 2010), as evidenced by the proteomics data. Interestingly, one lipid compound (Cer 18:0_18:0) which was significantly reduced in non-healthy lungs, has also been found dysregulated in the blow of a killer whale with bronchopneumonia (Harsla et al., 2024). Though this study only later diagnosed the bronchopneumonia in this animal, the similarity with our study may strengthen marker reliability to monitor health states. As surfactant is secreted by ATII cells in the lung, we cannot exclude measuring a proportion of it. Considering that marine mammals possess a surfactant composition optimised for an aquatic environment (Gutierrez et al., 2015; Miller et al., 2006; Spragg et al., 2004), and surfactant impairment has been linked with inflammatory lung diseases in humans (Griese, 1999), a combination of comparative lung tissue and surfactant analyses should be further explored to investigate potential pathological alterations.

Our results indicate a chronically inflamed lung in non-healthy harbour porpoises, along with implications of energetic imbalance and insufficient regenerative capacity. Taken together, this suggests progressive tissue loss and impairment of lung function, which may be exacerbated by a persistent cycle of sustained inflammation and inadequate repair.

### MultiOmics of the muscle

The muscle is a swiftly adaptable tissue, thus possesses a high turnover rate to build and maintain tissue (Frontera & Ochala, 2015; Howard et al., 2020). This may be reflected in the high elevation for various processes and features associated with muscle structure and function in non-healthy harbour porpoises, supporting our transcriptomic results from the same animals (Dönmez et al., 2026). However, regenerative processes are energetically costly. While proteome studies of Northern elephant seal muscles suggested a strictly controlled protein turnover during fasting periods (Khudyakov et al., 2022), this may not be true for harbour porpoises. Harbour porpoises are prone to starvation due to their restricted energy reserves, temperate-to-cold habitats and high metabolic field rate, resulting in the need to constantly forage to ensure insulation and energetic supply (Koopman et al., 2002; Rojano-Doñate et al., 2024). Additionally, a diseased condition as observed here, not only manifesting in local lung inflammation, but also in a potentially chronic tissue damage and systemic inflammation, increases energetic imbalance due to the persistent activation of repair and immune pathways. Upon energy deprivation and increasing overall ATP demand, muscle tissue will be utilised as primary metabolic reservoir to meet energetic requirements (Wolfe, 2006). In accordance with this, we found pathways and metabolites pointing to protein degradation and muscle atrophy. Similar findings have been observed in the muscles of chronic obstructive pulmonary disease (COPD) patients (Jagoe & Engelen, 2003) and in the plasma of diseased dolphins (Derous et al., 2022). A potentially impaired or altered energy metabolism in muscles of non-healthy harbour porpoises was further underlined by decreased fatty acid β-oxidation. Toothed whales feed exclusively on a high-fat diet, generate their energy primarily through lipid metabolism and have lost the ability for ketogenesis (Wolfgang et al., 2021). An impairment of their main energy pathway may force them to rely on alternative, less energetically favourable pathways such as muscle protein catabolism to provide amino acids for gluconeogenesis.

As post-mitotic tissue, muscles are prone to accumulation of oxidative injury (Frontera & Ochala, 2015; Rom & Reznick, 2016). To prevent exceeding oxidative stress, marine mammal muscles possess high baseline antioxidant concentrations (Allen & Vázquez-Medina, 2019; García-Castañeda et al., 2017; Vázquez-Medina et al., 2006). Likewise, we could identify multiple antioxidative features, including glutathione-associated proteins and selenoamino-involved metabolites. Selenoamino acids are linked to glutathione metabolism which is enhanced in marine mammals (Chung & Maines, 1981; Vázquez-Medina et al., 2006; Wilhelm Filho et al., 2002). Moreover, the high levels of anti-inflammatory and antioxidative plasmalogens in the muscle of non-healthy harbour porpoises indicate an enhanced oxidative stress state. Plasmalogens possess a unique vinylether-structure which enables them to bind free radicals (Brosche & Platt, 1998; Murphy, 2001) and reduced levels have been noted in inflammatory diseases (Paul et al., 2019). Although it is not possible to determine whether the increased immune and stress responses are primarily due to a possible reduction in oxygen supply caused by pathological lung lesions, chronic disease conditions, as observed in the animals studied here, will have an adverse effect on the entire body and organism.

Our findings suggest that, despite aquatic adaptations and respiratory specialisations, the persistent pulmonary immune activation contributes considerably to an elevated energy requirement with implications for the whole-body. The potential link between lung and muscle responses implies that chronic immune activation and impaired pulmonary repair mechanisms not only increase energetic costs, but may also impede oxygen availability for muscular energy generation, therefore accelerating muscle proteolysis to compensate for this systemic energy discrepancy.

### Overlapping features cross tissue

Across both tissues, we could identify several, similarly regulated features in each omics set, which may be of interest in regard to downstream effects from the lung to the muscle. Again, antioxidant and immune responses were among the most common associations in highly abundant cross-tissue regulated features (e.g., GSTP1, ferritin, immunoglobulins, histamine, methylhistamine, plasmalogens).

In both lung and muscle, the highest upregulated protein was Protein Dispatched Homolog 1 (DISP1). DISP1 is involved in hedgehog signalling, which has a role in the pathogenesis of COPD and pulmonary fibrosis, and has been proposed as COPD biomarker (Ancel et al., 2020; Cigna et al., 2012). The simultaneous upregulation of DISP1 in the pathological lung as well and corresponding muscle of non-healthy harbour porpoises may pose as valuable indicator of the systemic impact of a compromised respiratory tract.

Another interesting cross-tissue metabolite is isovalerylcarnitine, which was upregulated in both tissues of non-healthy harbour porpoises. Isovalerylcarnitine is produced during disrupted leucine catabolism, for which the significant decrease of leucine and isoleucine in the muscle of non-healthy harbour porpoises may be supporting evidence (McCalley et al., 2019; Tanaka et al., 1966). Isovalerylcarnitine can also be formed from carnitine and toxic isovaleric acid. Isovaleric acid is abundant in specialised, non-metabolised fat tissues (e.g., acoustic fats) and muscles of harbour porpoises (Koopman, 2018), though it has not yet been quantified in the lung (Arregui et al., 2020; Lovern, 1934). In recent studies of acute respiratory distress syndrome and pathogen-associated pneumonia in humans, isovalerylcarnitine displayed distinctive abundance profiles between conditions and has also shown potential to be predictive of disease progression (den Hartog et al., 2024; Suber et al., 2023). As the immune responses in harbour porpoises showed similarities to those of humans, despite inhabiting different habitats, isovalerylcarnitine may also be suitable as biomarker indicative of lung diseases in harbour porpoises.

Since these dysregulated features likely circulate between tissues via the bloodstream, incorporating analyses of corresponding blood samples in future studies may deepen our understanding of multi-tissue pathophysiology and help assess their potential as biomarkers for respiratory disease in cetaceans.

### Potential signature to estimate lung health and disease in marine mammals

Lastly, we aimed to identify a potential feature profile that could be utilised in a biomarker panel to monitor and estimate lung health and disease in marine mammals. We applied an integrative MultiOmics approach, including proteomics, metabolomics and lipidomics of non-healthy and healthy harbour porpoises. Integrative MultiOmics are a relatively novel method that can provide comprehensive insights into the complex systems biology by simultaneously analysing different datasets of the same individuals (Lee et al., 2019). With mixOmics “DIABLO”, we identified a signature of a total of 18 proteins, metabolites and lipids in the lung and in the muscles that may drive the pathogenesis of lung-associated lesions in harbour porpoises. Our MultiOmics model indicated a strong interaction between SLC25A4, O-phosphoethanolamine and TG O-16:0_16:0_20:4 in the lung (Fig. 5D, Tab. 5). SLC25A4 is located in the mitochondrial membrane where it mediates ADP:ATP transport for energy generation and fuel (Clémençon et al., 2013), while TG O- is associated with lipid metabolism, pointing to a molecular signature in altered metabolism. In mice lungs exposed to ozone, O-phosphoethanolamine was associated with reduced hypersensitivity (Smith et al., 2023), which may validate its pathological role in the lung lesion-associated phenotype studied here. In the muscle, a signature of five features was observed within the 18 proposed features. This included a positive correlation between SPEG, a protein crucial for muscle function and myotube function (Campbell et al., 2021; Hsieh et al., 2000), and BMP 18:1_22_6, a bis-(monoacylglycero)-phosphate which are involved in vesicle trafficking (Showalter et al., 2020). The negative correlation of SPEG and BMP 18:1_22_6 with pipecolic acid, which is dysregulated in the serum of smokers and associated with oxidative stress (Dalazen et al., 2014; Shen et al., 2023), implies a compensatory response to an enhanced redox imbalance. Moreover, pipecolic acid was positively correlated with two anti-inflammatory and antioxidative plasmalogens (PE P-18:0_18:1, PE P-17:0_22:6; (Brosche & Platt, 1998; Murphy, 2001; Paul et al., 2019). This interaction may indicate that elevated oxidative stress triggers a protective shift in plasmalogen metabolism, thus further supports a potentially coordinated pattern to redox stress in the muscle. Together, these features cover most of the observed dysregulated pathways we observed in the muscle of non-healthy harbour porpoises, namely elevated oxidative stress, tissue degradation and impaired regeneration. This may qualify these features as a combined panel of markers for lung-associated muscle impairments, thereby enabling the analysis of the pathological consequences on distant organs.

Integrative MultiOmics studies have been mostly applied in biomarker identification for various dispositions and diseases (Bowerman et al., 2020; Harriott et al., 2025; Ivanova et al., 2024; Konigsberg et al., 2021). Compared to these studies, our samples size is rather small and could be larger for more predictive power. Nonetheless, the identified MultiOmics signatures are a first step for biomarker identification in marine mammals and additional functional analyses will give deeper insights into the molecular mechanisms and phenotype (Lam et al., 2020).

## CONCLUSION

Our integrative MultiOmics study on the lungs and locomotor muscles of free-ranging harbour porpoises indicates a profound and sustained immune activation in non-healthy individuals. Unlike the attenuating Th2 inflammation that is typically associated with parasitic infestation, the response reflected a proinflammatory type 1 activation, accompanied by impaired lung tissue repair, muscle atrophy and overall energetic imbalance. This molecular and metabolic dysregulation suggests reduced tissue function, potentially with direct consequences on diving ability and survival. Both lungs and muscles exhibited similarly altered abundance of several proteins, metabolites and lipids, implying systemic impacts, possibly mediated through circulation. Using mixOmics DIABLO, we identified highly correlated sets of molecules in both tissues that may serve as a lung disease-associated marker panel to support health assessments in cetaceans. Our results demonstrate that respiratory disease in harbour porpoises extends beyond the lungs and can be evident in muscles, thus affecting overall fitness and further emphasising the importance of considering systemic effects when evaluating population health. Because anthropogenic impacts, such as underwater-radiated noise, pollution and climate change, increasingly influence susceptibility to disease and resilience in cetaceans, these insights deepen the understanding of how global environmental pressures alter health and survival in marine sentinel species like the harbour porpoise.

### Limitations

While decomposition has to be considered when working with necropsy tissue material, we tried to minimise potential degradation effects by following standardised best practice necropsy and sampling protocols and only considering harbour porpoises within decomposition code 2 (max. 24 h post-death). As omics are an emerging field in marine mammal science, few comparable studies are published to compare the impact of decomposition and handling on detection. One study on surfactant found little lipid composition change of post-mortem sampled marine mammals for up to two days (Gutierrez et al., 2015). We recommend that future studies should implement time-comparative experiments, where possible, to support recommendations for best practices in marine mammal omics which will certainly continue to grow. However, we were able to identify proteins, metabolites and lipids from two tissues of deceased, free-ranging harbour porpoises and compare two physiologically distinct states. We analysed the altered regulation of metabolic, stress and immune responses and provide valuable insights into the systemic implications of respiratory diseases in harbour porpoises.

Since some individuals were by-caught and non-healthy animals often suffered from several diseases, we cannot exclusively attribute the immune mobilisation in the lung and oxidative stress and metabolic response of the muscle solely to the observed lung lesions, as various diseases have an effect on multiple organs (Celli et al., 2018; Nauffal et al., 2024; Reynolds, 2002). Additionally, this study focuses on free-ranging animals whose life history (last food intake, short-term stress, recent disease recovery, and other factors) is not known but can influence molecular responses. Still, inflammation is a hallmark reaction across many pathological conditions that impacts signalling responses like interleukin or HIF1 signalling, alters metabolic pathways such as shifting to higher utilisation of anaerobic metabolism, and influences common mechanisms, e.g., tissue remodelling.

Lastly, our relatively small sample size may have an effect on the results, especially if interindividual variability plays a role. Future studies should aim for larger sample sizes for more coherent results, although this is a limiting factor in marine mammal science because these animals are elusive, difficult to sample, and often strictly protected.

## DATA AVAILABILITY STATEMENT

The data sets of this study can be found in the article’s Supplementary Material and in online repositories. The names of the repository and accession numbers are listed in the Supplementary Material (Supplementary Tab. 1).

## ACKNOWLEDGMENTS

The authors thank all individuals who helped to collect the carcasses, perform the necropsies and conduct further investigations.

We thank the Core Facility Mass Spectrometric Proteomics as part of the Technology Platform Mass Spectrometry at University of Hamburg and University Medical Center Hamburg-Eppendorf for support with mass spectrometric measurements and analysis, funded by the Deutsche Forschungsgemeinschaft (German Research Foundation) – 247354600.

## CRediT AUTHORSHIP CONTRIBUTION STATEMENT

EMD: Conceptualisation, Data Curation, Formal Analysis, Funding Acquisition, Investigation, Project Administration, Validation, Visualisation, Writing – Original Draft Preparation; BS: Data Curation, Formal Analysis, Investigation, Project Administration, Validation, Writing – Review & Editing; BD: Data Curation, Formal Analysis, Investigation, Project Administration, Validation, Writing – Review & Editing; PN: Data Curation, Formal Analysis, Investigation, Writing – Review & Editing; DD: Visualisation, Writing – Review & Editing; AF: Conceptualisation, Funding Acquisition, Supervision, Writing – Review & Editing; US: Conceptualisation, Data Curation, Funding Acquisition, Resources, Supervision, Writing – Review & Editing

## ETHICS STATEMENT

Ethical review and approval were not required for the study since the investigated animals were collected once they were dead by stranding or bycaught within the German stranding network. The German stranding network conducts work, such as collecting and holding carcasses and samples from European protected species, following appropriate licenses from the relevant authorities.

